# Mechanisms of carbon dioxide detection in the earthworm *Dendrobaena veneta*

**DOI:** 10.1101/2023.04.12.536649

**Authors:** E Jordan Smith, Jennifer L Ryan, Sofia A Lopresti, Dastan BS Haghnazari, Karleigh AS Anderson, Sarah J Lipson, Erik C Johnson, Wayne L Silver, Cecil J Saunders

**Author notes:** **Author contributions:** First authorship – ES, Senior authorship – CS. Corresponding author(s): CS,; ES. ES, KA, EJ, WS, and CS contributed to the conception and design of the study. ES developed the exudate assay and with JR, SL, and DH, conducted experiments with that assay. KA and CS optimized the RNA extraction protocol. KA, SL, and CJS organized and analyzed transcriptomic data. ES wrote the first draft of the manuscript with major input from CJS. All authors read and approved the submitted version.

## Abstract

Carbon dioxide (CO_2_) is a critical biological signal that is noxious to many animals at high concentrations. The earthworm *Dendrobaena veneta* lives in subterranean burrows containing high levels of CO_2_ and respires through its skin. Despite the ecological and agricultural importance of earthworms, relatively little is known about how they make decisions in their environment, including their response to elevated levels of CO_2_. To examine CO_2_ detection in this species, we designed the exudate assay, in which we placed an earthworm in a sealed container, exposed it to varying concentrations of CO_2_ for one minute, and recorded the amount of exudate secreted. Because earthworms excrete exudate in response to noxious stimuli, we hypothesized that the amount of exudate produced was proportional to the amount of irritation. We repeated these experiments after treatment with several blockers for molecules with potential involvement in CO_2_ detection, including carbonic anhydrases, guanylate cyclase, TRPA1, ASICs, and OTOP channels. We also confirmed the presence of homologous transcripts for each of these gene families in an epithelial transcriptome for *D. veneta*. Additionally, since organisms often detect CO_2_ levels indirectly by monitoring the conversion to carbonic acid (a weak acid), we used the exudate assay to evaluate aversion to additional weak acids (formic acid, acetic acid, and propionic acid). Earthworms excreted significantly more exudate in response to CO_2_ in a dosage-dependent manner, and this response was muted by the general carbonic anhydrase inhibitor acetazolamide, the carbonic anhydrase IX/XII inhibitor indisulam, the calcium channel blocker ruthenium red, the sodium channel blocker amiloride, and the acid-sensing ion channel blocker diminazene aceturate. These data provide evidence of the role of carbonic anhydrase and epithelial sodium channels in earthworm CO_2_ detection, establish that, similar to other subterranean-dwelling animals, earthworms are extremely tolerant of CO_2,_ and contribute to our understanding of the mechanisms used by earthworms to detect and react to weak acids in their environment.

**Contribution to the field statement:** Carbon dioxide (CO_2_) is a major byproduct of cellular respiration and an important biological signal. The metabolism of animals living underground in burrows can cause the concentration of CO_2_ to increase above atmospheric CO_2_. Most animals have multiple molecular mechanisms that detect CO_2_ and find high concentrations aversive or noxious; some subterranean animals have adaptations that make them more tolerant of concentrated CO_2_. Earthworms live in burrows, are a keystone species for subterranean ecosystems, and in their lightless environment must rely primarily on tactile and chemical signals. Despite the profound importance of earthworms, few if any of the molecular mechanisms they use to detect these signals have been characterized. Here we use RNA sequencing to develop a list of candidate mechanisms that the European nightcrawler may use to respond to CO_2_. We also have developed an assay for comparing how noxious different concentrations of volatile compounds are to earthworms. In the present study, we utilize that assay with inhibitors to further narrow the possible multiple molecular mechanisms responsible for the detection of CO_2_. We suspect that our study is among the first attempts to combine RNA-sequencing technology and pharmacology to describe earthworm sensory biology.

## Introduction

As a major byproduct of cellular respiration, CO_2_ is pervasive in most ecosystems and may indicate the presence of other living organisms. It is thus a critical signaling molecule that is often attractive or aversive depending on the organism and concentration. Fruit flies, for example, avoid CO_2_ released by neighboring stressed flies (Suh et al., 2004), while mosquitoes are attracted to CO_2_ emitted by their hosts (Spanoudis et al., 2020). At higher concentrations, CO_2_ can be noxious, and may result in hypercapnia, hypoxia, or anesthesia (Cummins et al., 2019). Given its biological prevalence, potential health risks, and environmental relevance, it is critical to understand the mechanisms by which organisms detect CO_2_.

Important decomposers in some ecosystems and invasive species in others, earthworms play critical roles in environmental health and agriculture. In many ecosystems, earthworms are a key component of soil fertility, influencing soil turnover, soil aeration, and nutrient availability (Edwards, 2004). Despite their essential environmental role, we know little about what chemicals attract and repel earthworms nor the mechanisms by which they detect those chemicals (Silver et al., 2019; Reed et al., 2021). One such chemical is carbon dioxide (CO_2_). The current concentration of CO_2_ in air is approximately 0.04% (Amundson & Davidson, 1990; Scott, 2011). Earthworms live in burrows up to three meters deep and may encounter CO_2_ concentrations of 0.04% to 13.0% (Amundson & Davidson, 1990). Some subterranean organisms, such as naked mole rats, have unique adaptations that allow them to tolerate normally noxious concentrations of CO_2_ (Shams et al., 2005; Fang et al., 2014); earthworms may have similar adaptations.

The ubiquitous presence and broad importance of CO_2_ has produced multiple detection pathways across organisms (Figure 1A). Most mechanisms require carbonic anhydrase, which catalyzes the reversible conversion of carbon dioxide into a bicarbonate ion and a proton, reacting with water from extracellular fluid (Lindskog, 1997; Figure 1A). Carbonic anhydrase is found across animals, plants, and microorganisms, includes three independently evolved isozyme families, and is essential in many biological functions including metabolism, cellular transport, and acid-base balance (Henry, 1996; Banerjee & Deshpande, 2016). The multiple isoforms of α-carbonic anhydrases, the family found in animals, are labeled CAI to CAXV and are differentially expressed among tissues and cell types (Tarun et al., 2003). Mechanisms of CO_2_ detection may respond to CO_2_ directly, to bicarbonate ions, or to protons from the carbonic anhydrase reaction (Makino et al., 2019; Figure 1A).

**Figure 1.**
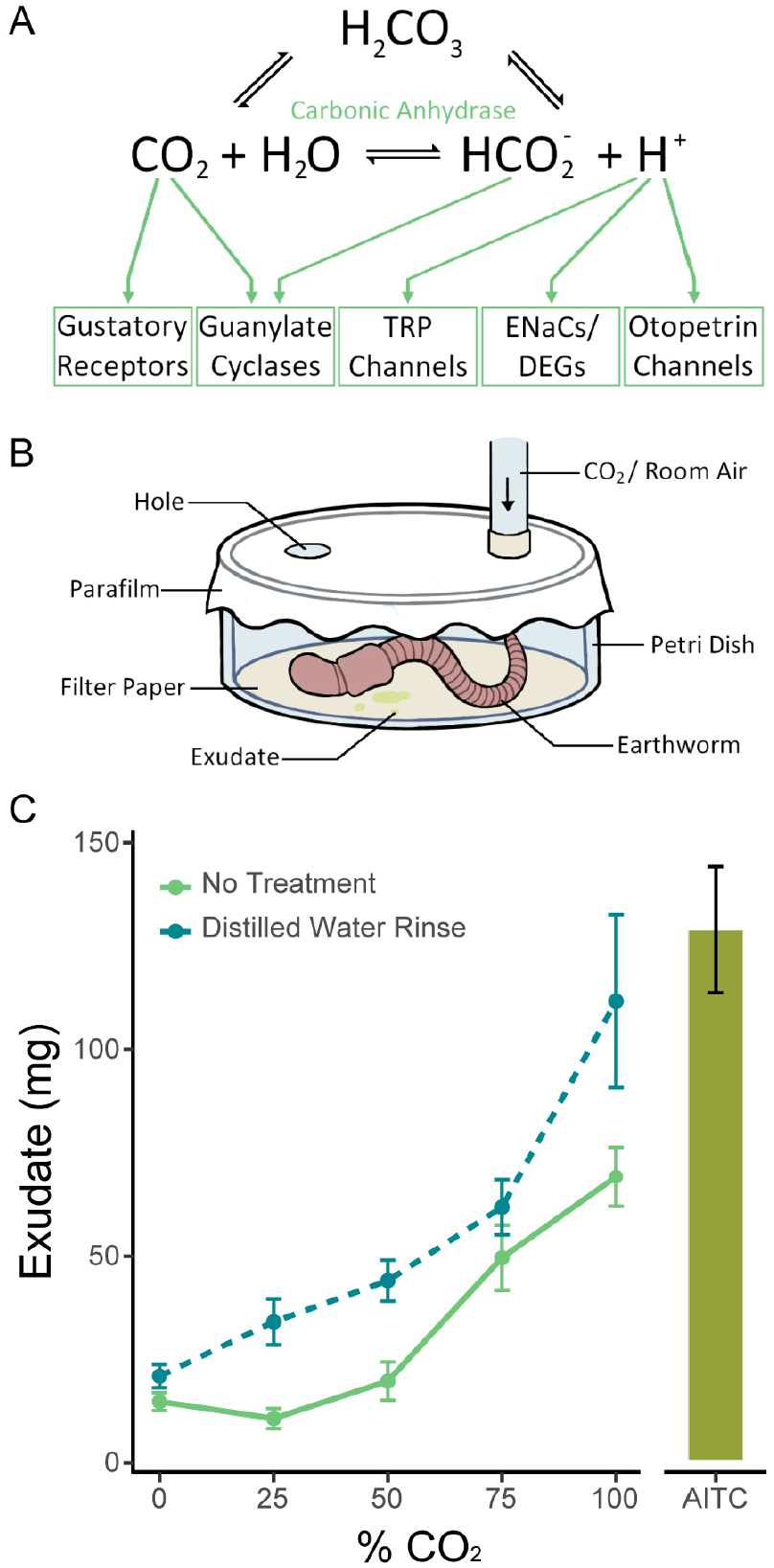
Mechanism of CO_2_ detection and the exudate assay. (A) The chemical reaction CO_2_ + H_2_O ←→ HCO_2_^-^ + H^+^ is catalyzed by carbonic anhydrase. The green arrows point to the potential mechanisms by which CO2 and its products could be detected. (B) Diagram of the exudate assay. Earthworms were placed in a sealed container filled with a volatile or gaseous chemical for one minute, and the amount of exudate produced in that time was recorded. (C) Earthworms excreted significantly more exudate in response to increasing concentrations of CO_2_ (p<0.0001, n=12-20 one-way ANOVA). Earthworms also excreted significantly more exudate in response to CO_2_ if they were rinsed in deionized water prior to the assay (p<0.0001, two-way ANOVA). The response to air flowed over filter paper containing 500 µl of 500 mM AITC, a known noxious stimulus, is shown for the purposes of assay validation. Graphed values are means ± SEM.

*D. veneta* lives in burrows several cm below the ground and detects chemicals in the soil using sensory cells on its epithelium, either grouped into epithelial sensory organs (ESOs) or found alone as solitary chemoreceptor cells (Hess, 1925; Csoknya et al., 2005; Kizsler et al., 2012). The cells project to the ventral nerve cord and are categorized into five types based on fine structure (Csoknya et al., 2005; Kizsler et al., 2012). These organs are more abundant on the anterior segments and are the likely candidates for the cellular receptors responsible for CO_2_ and weak acid detection. However, this study is the first attempt to determine the molecular identity of the genes responsible for CO_2_ detection in these tissues.

Here we investigate CO_2_ and weak acid detection in the European Nightcrawler *Dendrobaena veneta* (previously known as *Eisenia hortensis*) using known mechanisms from other species as a template. Ionotropic gustatory receptors (GRs), acid-sensing ion channels (ASICs), guanylate cyclases (GCs), otopetrin channels, and transient receptor potential ankyrin 1 (TRPA1) channels have all been implicated in the detection of CO_2_ or weak acids like carbonic acid in other animals. There is some evidence of these mechanisms in *D. veneta*. Earthworms likely have functioning TRPA1 channels, as they show behavioral aversion to TRPA1 activators (Silver et al., 2018). We expected high concentrations of CO_2_ to be noxious to earthworms as in other species, and we hypothesized that carbonic anhydrases, guanylate cyclases, TRPA1, ENaCs, and OTOPs may be involved with CO_2_ detection in earthworms based on the analysis of a *D. veneta* epithelial transcriptome presented in this study.

## Materials and Methods

### Earthworms

*D. veneta* earthworms were housed in plastic bins (approximate volume 40L) filled half-way with moist topsoil covered with paper. The bins were maintained at room temperature with two light sources placed above the bins on a 12-12 light-dark cycle. The worms were watered with 250 ml of water poured over the newspaper twice a week and fed with approximately 5 g of “Purina Worm Chow” once a week. Only adult earthworms, identified by their visible clitellum, were used in experiments.

### CO_2_ Aversion

Earthworms excrete exudate in response to noxious stimuli (Heredia et al., 2008). Given that high concentrations of CO_2_ are aversive in most species, we predicted that the degree of exudate production would reflect CO_2_ detection and relative aversiveness. To perform the exudate assay, we first gently rinsed the earthworms in tap water and placed them in glass containers with moist paper towels to deprive them of soil and allow their digestive tract to empty. After 24 hours, a worm was placed in a glass petri dish with filter paper at the bottom and folded up the sides, and the dish was sealed with parafilm. The parafilm was punctured twice, and the dish was then saturated with air containing some percentage of CO_2_ through a nozzle inserted through the parafilm for one minute. The filter paper was weighed before and immediately after each experiment to calculate the weight of exudate excreted (Figure 1B).

Using this method, we tested responses to atmospheric CO_2_, 25% CO_2_, 50% CO_2_, 75% CO_2_, and 100% CO_2_ and analyzed the data with a one-way ANOVA and a TukeysHSD. Varying percentages of CO_2_ were obtained by combining different flow rates of room air and 100% CO2 using rotameters and a bubble flow meter for a final flow rate of approximately 32 ml/s for all trials. In this assay, 0% CO_2_ represents room air, which typically has 400-1,000 ppm CO_2_ (approximately 0.04%), 25% CO_2_ includes approximately 250,525 ppm CO_2_, 50% includes approximately 500,350 ppm CO_2_, 75% includes approximately 750,175 ppm CO_2_, and 100% includes approximately 1,000,000 ppm CO_2_. These results were compared to treatment with distilled water using a two-way ANOVA and TukeysHSD. All exudate assay data was analyzed and graphed in R (R Core Team, 2021) using the packages tidyverse (Wickham et al., 2019), dplyr (Wickham et al., 2023), and ggplot2 (Wickham, 2016). Figures were then manually edited to improve readability and aesthetics. Relevant code for reproducing these methods is available on Github and data files are available with this publication.

### Transcriptomics

The epithelium from the prostomium (1^st^ segment) and 15^th^ segment posterior to the clitellum (hereafter referred to as the mid-segment) were quickly dissected free from the underlying muscle and flash frozen on glass depression slides placed on dry ice. Once frozen solid, the tissue was transferred into 1.5 mL LoBind microcentrifuge tubes (Eppendorf) where it was crushed with a plastic pestle; both items were maintained on dry ice prior to use to maintain their freezing temperature during this process. Refrigerated TRIzol Solution (1mL, Thermo Fisher Scientific) was added, and tissue was further macerated until no intact fragments were visible. Subsequently, RNA was isolated from the tissue-TRIzol mixture using the manufacturer’s standard chloroform extraction and isopropanol precipitation protocol. RNA concentration and integrity was verified using a bioanalyzer (Agilent). This protocol was developed because the earthworm epithelium was surprisingly durable, and other standard methods for tissue disruption (e.g. tissue homogenizers and column systems) failed to yield enough mRNA for sequencing. We dissected the prostomium and mid-segments from 9 earthworms in total. We combined tissue from 3 individuals before extracting RNA to give us 3 prostomium samples and 3 mid-segment samples.

RNA libraries were prepared from total RNA using the Kapa Stranded mRNA-Seq library prep kit, and 150 bp paired-end sequencing was performed on an Illumina HiSeq 4000 at Duke University’s Center for Genomic and Computational Biology (Durham, NC). Trimmomatic v0.36 (Bolger et al, 2014) was used to filter raw reads and to remove Illumina adaptors (4bp, mean Q30). *De novo* transcriptome assembly was performed by processing the filtered reads with Trinity v.2.5.1 (Haas et al., 2013), and the resulting transcripts were annotated with Trinotate v.3.2.2 (Bryant et al., 2017) using the recommended settings. To develop gene candidate lists, the resulting Trinotate database was filtered for transcripts that were predicted to code for protein and contained one of the following terms: “carbonic anhydrase,” “gustatory receptor” “acid sensing ion channel,” “ASIC,” “guanylate cyclase,” “otopetrin,” “transient receptor potential,” or “TRPA.” These results were then confirmed and collected into candidate lists via human curation and are provided as used for analysis in Supplemental Material 1. Cladograms depicting the similarity of the predicted protein sequence for each group of candidate transcripts were constructed with R (R Core Team, 2021) using the following packages: ggtree (Yu et al, 2017), msa (Bodenhofer et al., 2015), seqinr (Charif et al., 2007), tidyverse (Wickham et al., 2019). We also conducted a differential expression analysis between RNA extracted from the prostomium and midsegment epithelium using DESeq2 Bioconductor package (Love et al., 2014). Relevant code for reproducing these methods is available on Github and sequencing data has been uploaded to the NCBI sequence read archive.

### Pharmacology

We repeated the exudate assay after treatment with the inhibitors and blockers described in Table 1. Inhibitors and blockers were sourced from Tocris Bioscience (Acetazolamide, Diminazene Aceturate, S4, Topiramate) and Sigma-Aldrich (Amiloride, HC030013, Methylene Blue, Ruthenium Red, ZnCl_2_, U-104). For these treatments, the earthworms were placed in a 10 ml conical tube filled with 5 ml of the blocker prior to the experiment. After 10 minutes, they were removed from the tube and gently dried with a paper towel before being placed in the assay chamber. The TRPA1 activator AITC, which we have previously shown to be aversive in *D. veneta* (Smith, 2019) and *Lumbricus terrestris* (Silver et al., 2019), was used as a positive control. In these trials, 500 ul of 500 mM AITC (allyl isothiocyanate) was pipetted onto a strip of filter paper placed in a 10 ml pipette, and air was flown through the pipette into the assay chamber. The data were analyzed with two-way ANOVAs and TukeysHSDs. p-values were adjusted using a Bonferroni correction for the carbonic anhydrase inhibitors, the receptor blockers, and the AITC controls. We observed no mortality during exudate assay experiments, but we did notice less movement or activity after some treatments; AITC controls allowed us to assess whether treatments with significant results affected exudate production rather than CO_2_ detection.

**Table 1.**
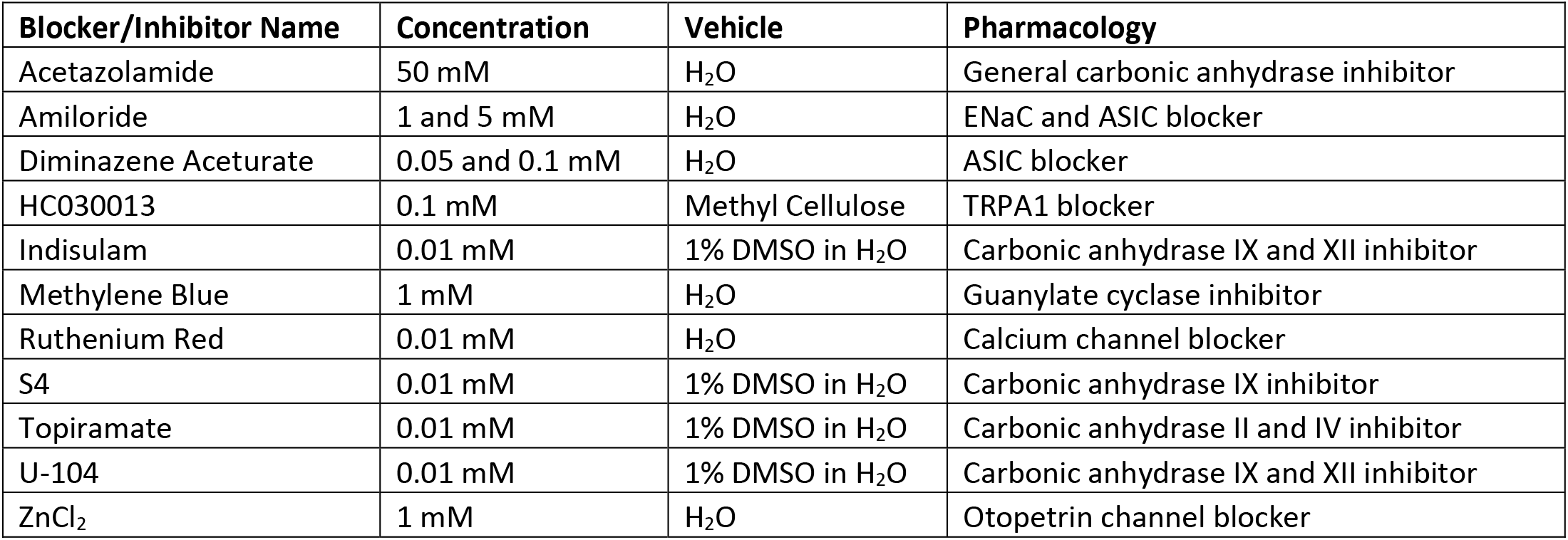
Blockers used in the exudate assay with their concentrations, vehicles, and pharmacological interactions of interest.

### Organic Acids

CO_2_ detection often relies on the detection of a weak acid (carbonic acid). Therefore, we also studied the responses of earthworms to weak organic acids to assess potential overlapping mechanisms. As detritivores, earthworms likely encounter fermented organic matter that contains organic acids in nature. Formic acid, acetic acid, and propionic acid (Fisher Scientific) were tested at concentrations of 50, 100, 250, 500, 1000 ppb in the exudate assay. For these trials, 250 ul of acid was pipetted onto a strip of filter paper placed in a 10 ml pipette, and air was flown through the pipette into the assay chamber to generate vapor. These trials were repeated at 250 ppb after exposure to the sodium channel and ASIC blocker amiloride using the procedure described above. These data were analyzed using two-way ANOVAs and TukeysHSDs.

## Results

### CO_2_ Aversion

Earthworms excreted the most exudate in response to 100% CO_2_. Over the course of the one-minute assay, earthworms first excreted clear liquid, and then a thicker pale green substance. Levels of movement varied between earthworms. Earthworms excreted more exudate with increasing concentrations of CO_2_ in a dosage-dependent manner (p<0.0001, n=12-20, two-way ANOVA), where concentrations of 0%, 25%, 50%, 75%, and 100% were tested (Figure 1C). Compared to these results, significantly more exudate was excreted when the earthworms were rinsed with deionized water before the experiment (p<0.0001, n=6-12, two-way ANOVA). Washing the worms prior to the experiment likely removes mucus and epithelial surface liquid that has a greater buffering capacity than deionized water, making the earthworm’s epithelium more sensitive to changes in pH.

### Transcriptomics

The assembled *D. veneta* epithelial transcriptome contained 36 protein coding transcripts annotated as carbonic anhydrase (CA) homologs (Figure 2). The presence of CA mRNA supports the idea that these enzymes are involved in an earthworms ability to detect CO_2_ concentration. Particularly of note are homologs to CAVI, one of the only known secreted CA isoforms which is found in saliva and respiratory epithelium surface liquids of many other species (Leinonen et al., 2004; Fábián et al., 2015). We also queried our transcriptome for genes known to be involved in the detection of CO_2_ and the various ions associated with carbonic acid and found many.

**Figure 2.**
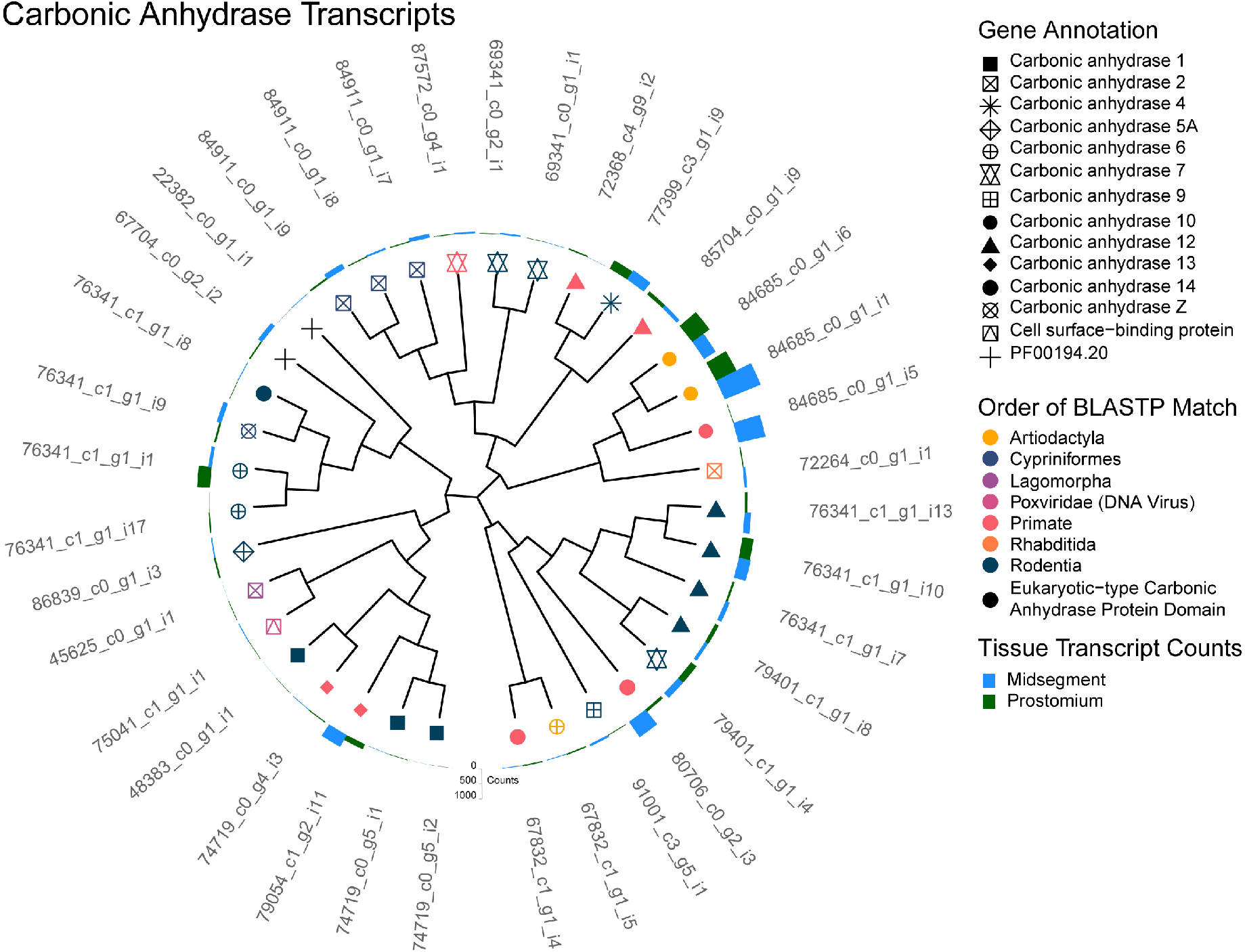
Carbonic Anhydrase Transcripts. A cladogram depicting the predicted protein sequence similarity between unique transcripts that annotated as carbonic anhydrase (CA). Predicted CA homologs identified by BLASTP are represented as the symbol shape and the order of the homolog as the symbol color. Note that transcript 75041_c1_g1_i1 (pink triangle inside square) showed homology with a viral cell surface-binding protein that contains a protein domain similar to eukaryotic-type carbonic anhydrase. Additionally, two transcripts (black plus sign; 22382_c0_g1_i1 and 67704_c0_g2_i2), did not have a BLASTP homolog but were annotated by Pfam with a eukaryotic-type carbonic anhydrase protein domain (PF00194.20). Total transcript counts from prostomium (green) and midsegment (blue) RNA are depicted as bars at the end of each tree tip.

We did not find any transcripts, however, which annotated as homologs of the ionotropic gustatory receptor family. In insects and many other invertebrates, CO_2_ directly activates gustatory receptors, a family required for taste and pheromone detection (Kwon et al., 2007; Ning et al., 2016; Chu et al., 2020). *C. elegans* also detects CO_2_ directly with isolated chemosensory BAG neurons (Smith et al., 2013). To ensure that our filtering criteria were not artifactually eliminating this gene family, we filtered our annotated transcriptome for any occurrence of the term “gustatory” but did not prefilter for protein-coding transcripts. We still found no transcripts annotated as belonging to this gene family. Automated gene annotation pipelines can undercount ionotropic sensory receptors (Agnihotri et al., 2016; McKenzie et al., 2016; McKenzie & Kronauer, 2018), but it would be uncommon for there to be no evidence of transcripts corresponding to ionotropic gustatory receptors if they were enriched in the epithelial tissues we sequenced.

In contrast to the absence of gustatory receptors, 29 unique transcripts annotated as guanylate cyclase were identified in the epithelial transcriptome (Figure 4). Bicarbonate ions sometimes activate guanylate cyclases, opening cGMP-sensitive ion channels and increasing cGMP (Sun et al., 2009). This is true in mice, where receptor-type guanylate cyclases GC-G and GC-D are activated by low concentrations of CO_2_ (Kuhn, 2016), as well as in *C. elegans*, where chemosensory BAG neurons require the receptor-type guanylate cyclase GCY-9 for their response (Hallem et al., 2011). GCs are more commonly considered to be essential proteins in G-protein-coupled receptor signaling cascades than as direct detectors of CO_2_ or bicarbonate. However, their presence in the epithelial transcriptome affirms this gene family’s inclusion on our candidate list.

Of the candidate detectors that respond to hydrogen ions produced by the carbonic anhydrase reaction directly, ASICs were the most abundant in the transcriptome with 60 unique transcripts annotated (Figure 3). ASIC channels belong to the Epithelial Na^+^ channel (ENaC)/degenerin (DEG) channel superfamily. In invertebrates, these channels have been implicated in salt detection, sodium absorption, blood pressure regulation, mechano-transduction, and acid-sensing (Kellenberger & Schild, 2002; Carattino & Montalbetti, 2020). ASIC homologues have been characterized in *C. elegans* (Rhoades et al, 2019), and there are many DEGs and ENaCs found across invertebrates (Hanukoglu & Hanukoglu, 2016). While ASIC blockage muted CO_2_ responses in rats (Akiba et al., 2007); the subtype ASIC3 is involved in chemoreception and opens at the physiologically relevant pHs of 7.3 to 6.7 (Osmakov et al., 2014; Li Xu, 2011), and could thus be involved in CO_2_ detection. However, loss of function in mice did fully ablate CO_2_ responses in these animals (Detwieler et al., 2018). One of the 60 ASIC transcripts is annotated as an ASIC3 homolog (Figure 3, S1, 87824_c0_g1_i27). However, the relatively low percent sequence identity (26.5 - 50%, S1) between these transcripts and their ASIC homologs makes us hesitant to narrow this candidate list further based on the Trinotate annotations alone.

**Figure 3.**
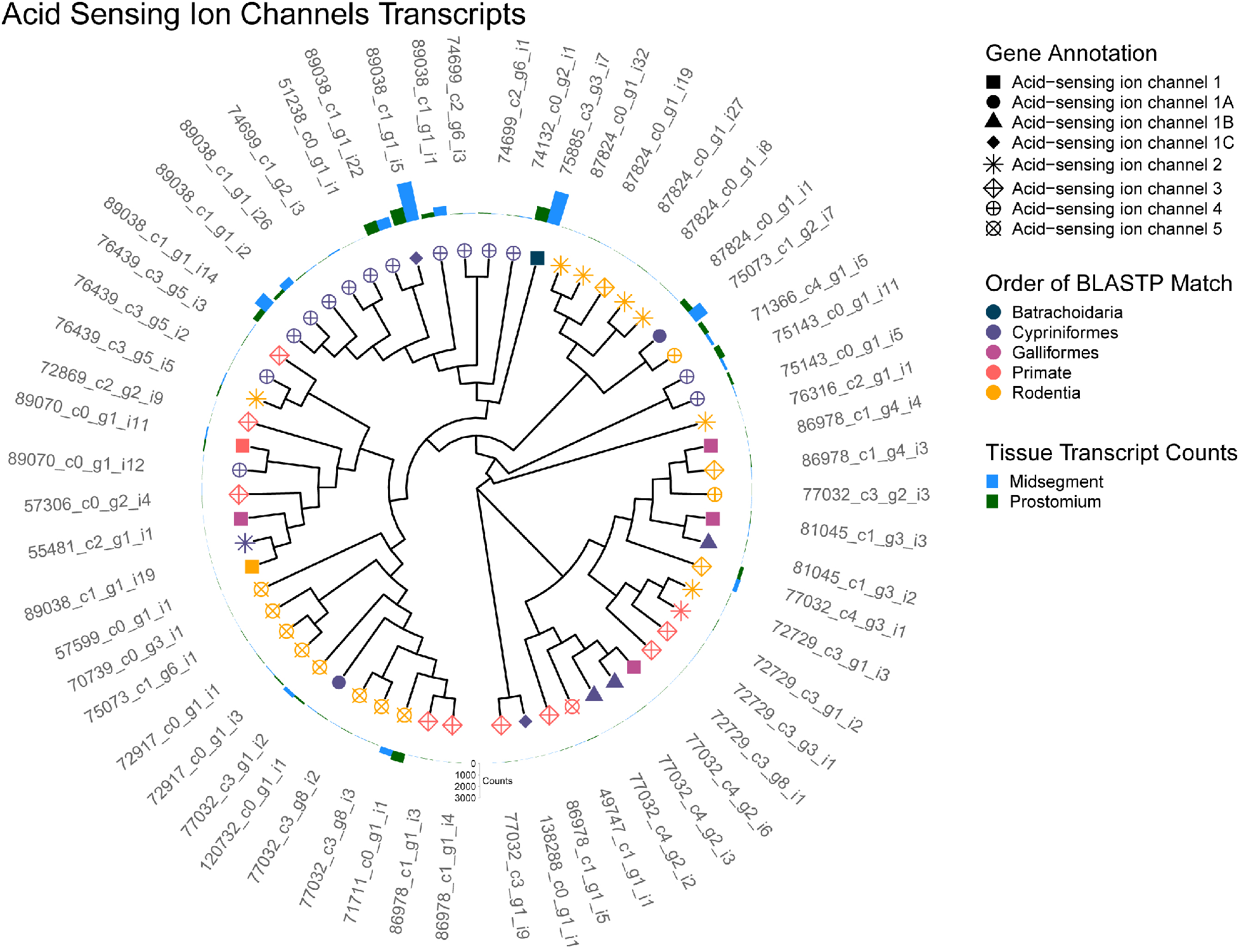
Acid Sensing Ion Channel (ASIC) Transcripts. A cladogram depicting the predicted protein sequence similarity between unique transcripts that annotated as ASICs. Predicted ASIC homologs identified by BLASTP are represented as the symbol shape and the order of the homolog as the symbol color. Total transcript counts from prostomium (green) and midsegment (blue) RNA are depicted as bars at the end of each tree tip.

**Figure 4.**
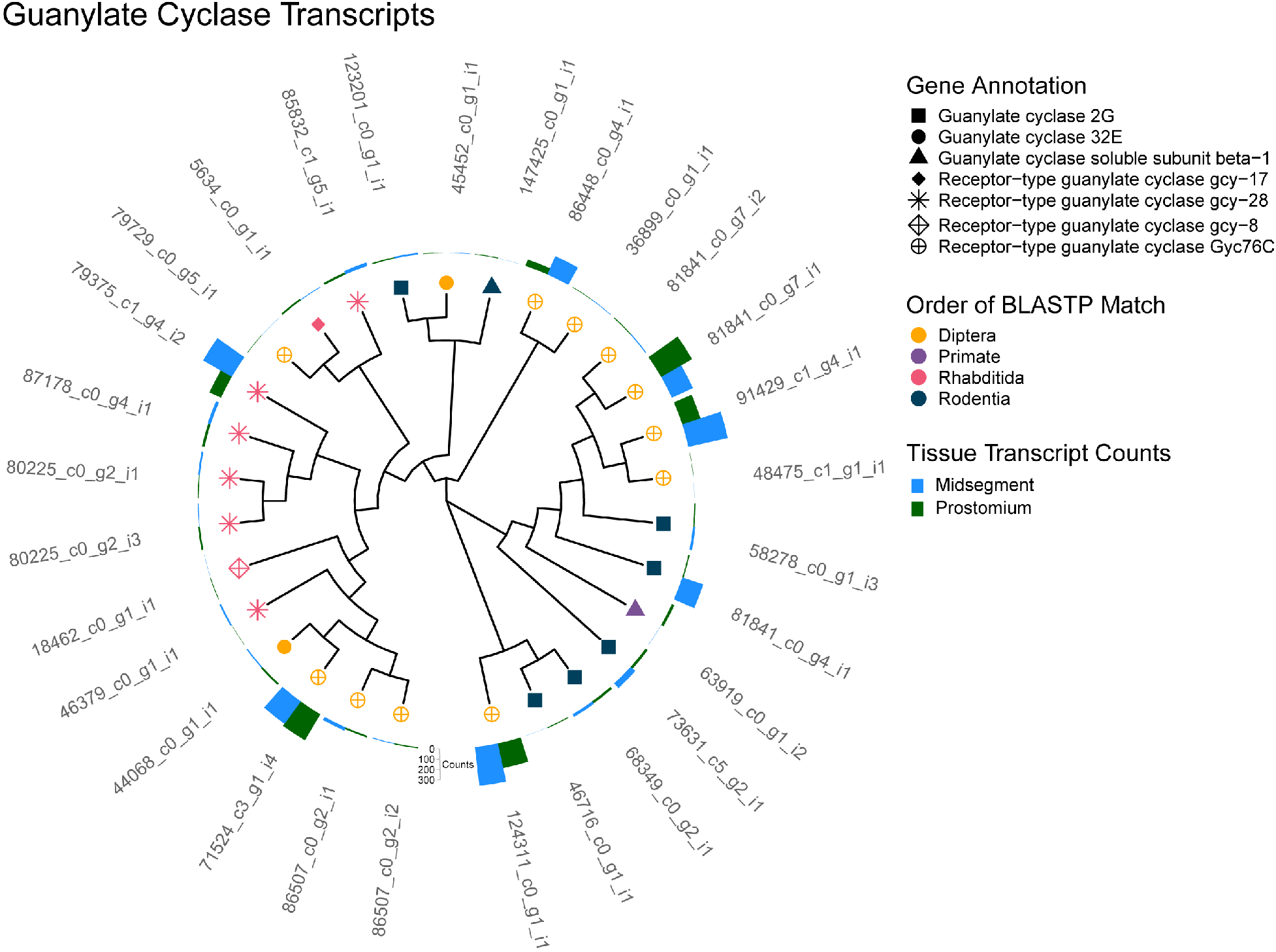
Guanylate Cyclase (GC) Transcripts. A cladogram depicting the predicted protein sequence similarity between unique transcripts that annotated as GC. Predicted GC homologs identified by BLASTP are represented as the symbol shape and the order of the homolog as the symbol color. Total transcript counts from prostomium (green) and midsegment (blue) RNA are depicted as bars at the end of each tree tip.

Analysis of the earthworm epithelial transcriptome returned 20 transcripts annotated with an otopetrin protein domain (PF03189.12), and 11 of these were identified as homologs of *Drosophila* otopetrin (OTOP) via BLASTP (Figure 5, S1). OTOP1 and OTOP3 are conserved across species with related genes in *Drosophila* (Tu et al., 2018). OTOP1, the proton-sensitive proton channel required for sour taste in mammals (Teng et al, 2019; Tu et al, 2018), may also respond to weak acids such as carbonic acid, a product of the carbonic anhydrase reaction (Teng et al., 2019).

**Figure 5.**
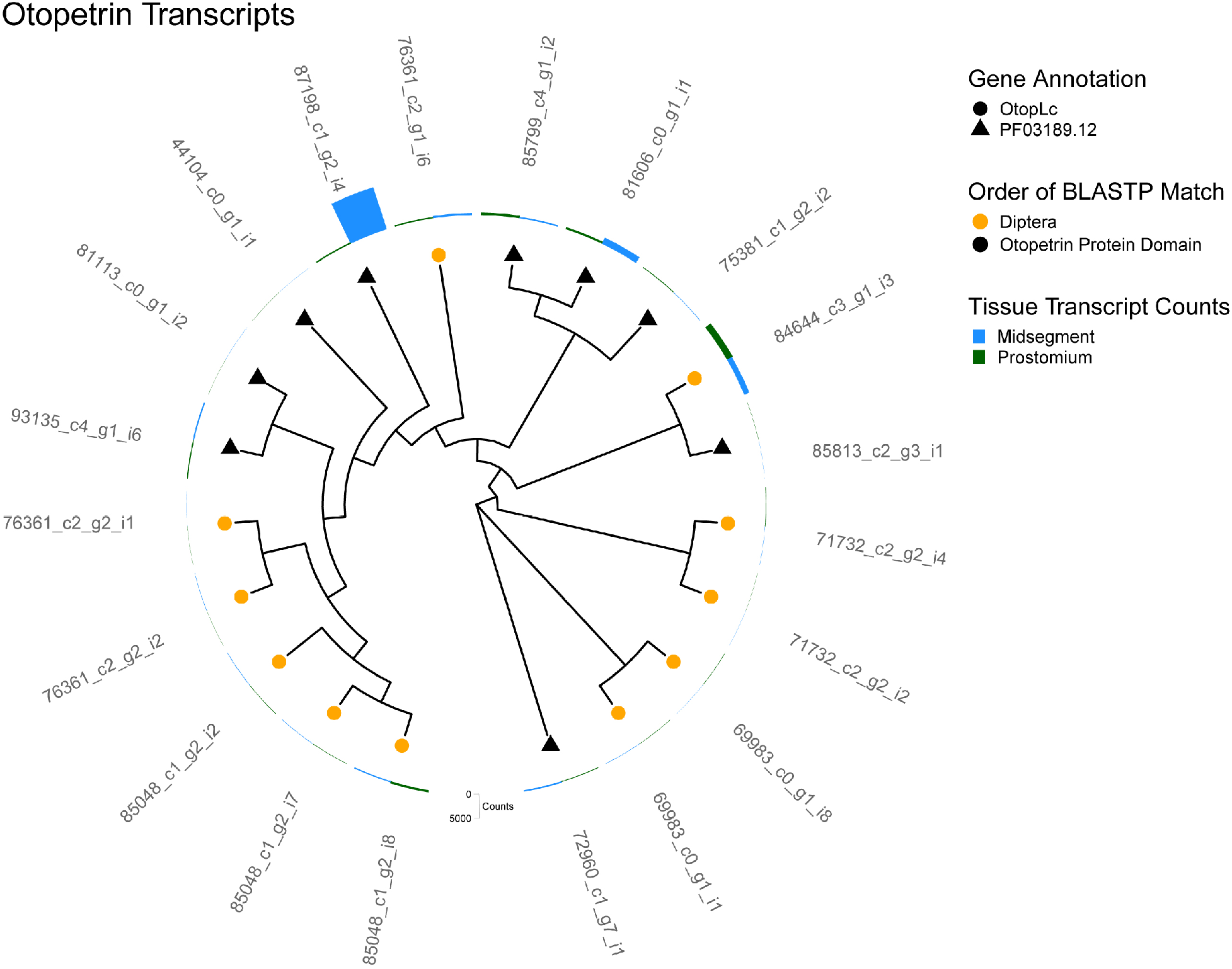
Otopetrin Transcripts. A cladogram depicting the predicted protein sequence similarity between unique transcripts that annotated as an otopetrin. Predicted otopetrin homologs identified by BLASTP are represented as the symbol shape and the order of the homolog as the symbol color. Additionally, nine transcripts (black triangle), did not have a BLASTP homolog but were annotated by Pfam with a otopetrin protein domain (PF03189.12). Total transcript counts from prostomium (green) and midsegment (blue) RNA are depicted as bars at the end of each tree tip.

TRPA1 is a calcium channel conserved throughout metazoan life that reacts to irritants, endogenous inflammatory agents, temperature, and weak acids (Bandell et al., 2004; Bautista et al., 2006; Wang et al., 2011). We found 9 transcripts in the earthworm transcriptome that shared homology with TRPA1 (Figure 6). There is evidence in mice and *in vitro* that the response of sensory neurons to CO_2_ is mediated by TRPA1 channels activated by intracellular acidification from protons (Takahashi et al., 2008; Wang et al., 2010; Wang et al., 2011). Furthermore, many of the vermifuges used to collect earthworms from the soil are TRPA1 agonists, meaning that there is much behavioral evidence that suggests earthworms have functional TRPA1 channels (Silver et al., 2019).

**Figure 6.**
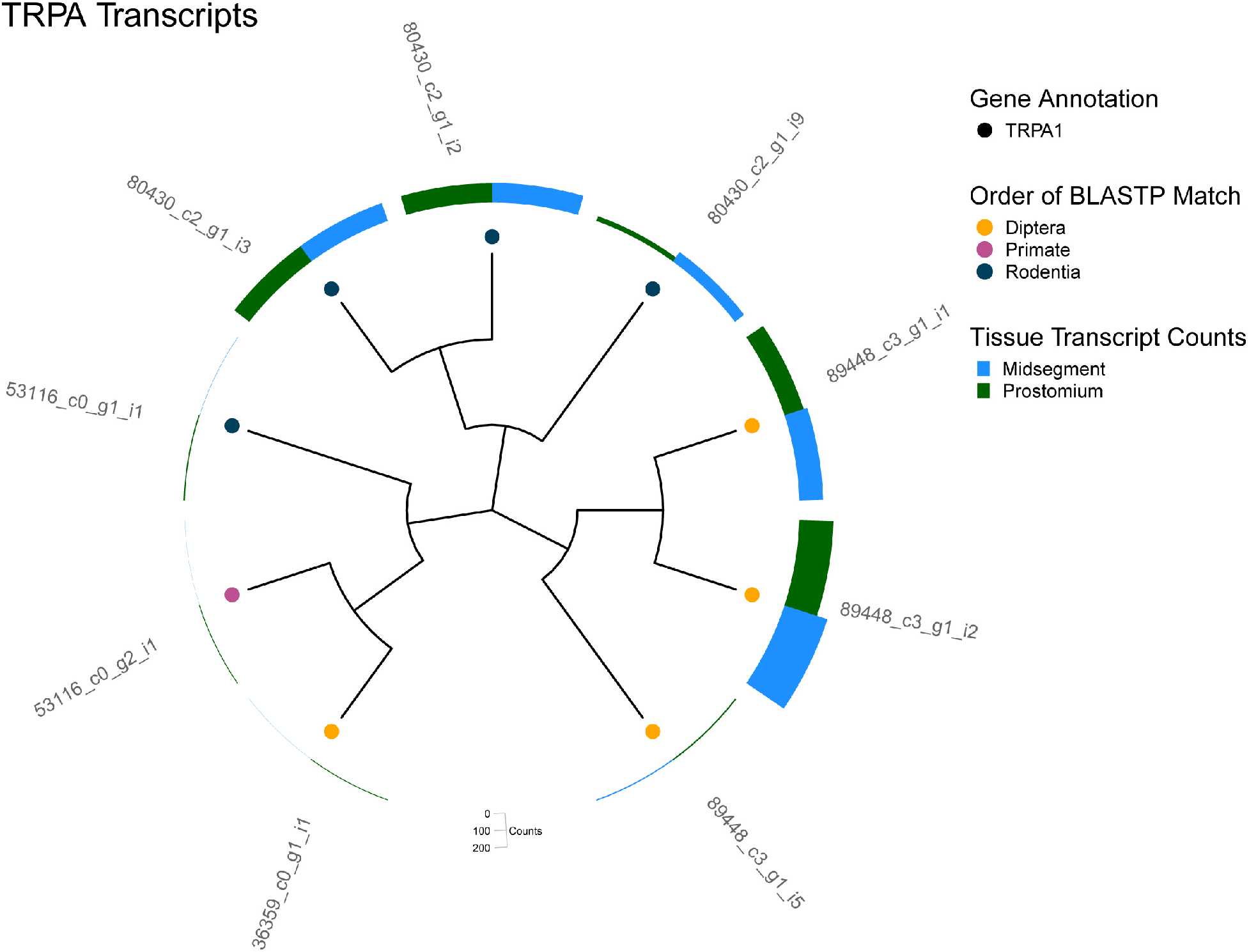
TRPA Transcripts. A cladogram depicting the predicted protein sequence similarity between unique transcripts that annotated as TRPA1. Predicted TRPA1 homologs identified by BLASTP are represented as the symbol shape and the order of the homolog as the symbol color. Total transcript counts from prostomium (green) and midsegment (blue) RNA are depicted as bars at the end of each tree tip.

In addition to using a *D. veneta* epithelial transcriptome to generate a list of candidate genes, we also conducted differential expression analysis between two epithelial tissues. While purported sensory cells are found along the entire length of the earthworm, these cells are more concentrated on the first segments than on segments towards the middle of the body (Csoknya et al., 2005). Since these cells are more abundant near the animal’s head, we hypothesized that transcripts more enriched in this tissue would be more likely to have a sensory function and might provide a clue to the genes responsible for CO2 detection. Transcript counts in both tissues are depicted as bars at the end of each tree branch for CA (Figure 2), GC (Figure 3), ASICs (Figure 4), OTOP (Figure 5), and TRPA1 (Figure 6). While this analysis detected 67 transcripts that were significantly differential expressed (adjusted p < 0.05) between the two tissues, none of these transcripts were annotated as genes involved in the detection of CO_2_ or related chemosensory processes (S2). Thus, we chose to test if any of the genes in our candidate list might be involved in the earthworm’s response to noxious levels of CO_2_.

### Pharmacology

The exudate assay was repeated with 0%, 50%, and 100% CO_2_ after treatment with carbonic anhydrase inhibitors and blockers for receptors potentially involved in CO_2_ detection (Table 1). As rinsing the earthworms in water increased their sensitivity to CO_2_ (Figure 1C), all blocker trials were compared to trials with the blocker’s respective vehicle (water, methyl cellulose, or 1% DMSO).

The broad-spectrum CA inhibitor acetazolamide significantly muted the response to CO_2_ (Figure 7A, p<0.01, n=8-13). This data supports the hypothesis that earthworms are responding to the presence of bicarbonate or hydrogen ions rather than detecting CO_2_ directly. Attempts to utilize selective CA inhibitors to determine the major CA isoforms involved in the conversion of CO_2_ were less clear. Indisulam, a blocker of CA IX and CA XII (p<0.05, n=6-8), significantly muted the response to CO_2_ (p>0.05, n=6-8), but U-104, which also blocks CA IV and CA XII, and S4, which also blocks CA IX, resulted in no significant changes. Topiramate, which blocks CA II and CA IV, also did not significantly mute the response to CO2 (p>0.05, n=6-8). The most parsimonious explanation of these results is that one or more isoforms of earthworm CA IX and CA XII have sufficiently diverged from isoforms in other species to make U-104 and S4 ineffective blockers in earthworms.

**Figure 7.**
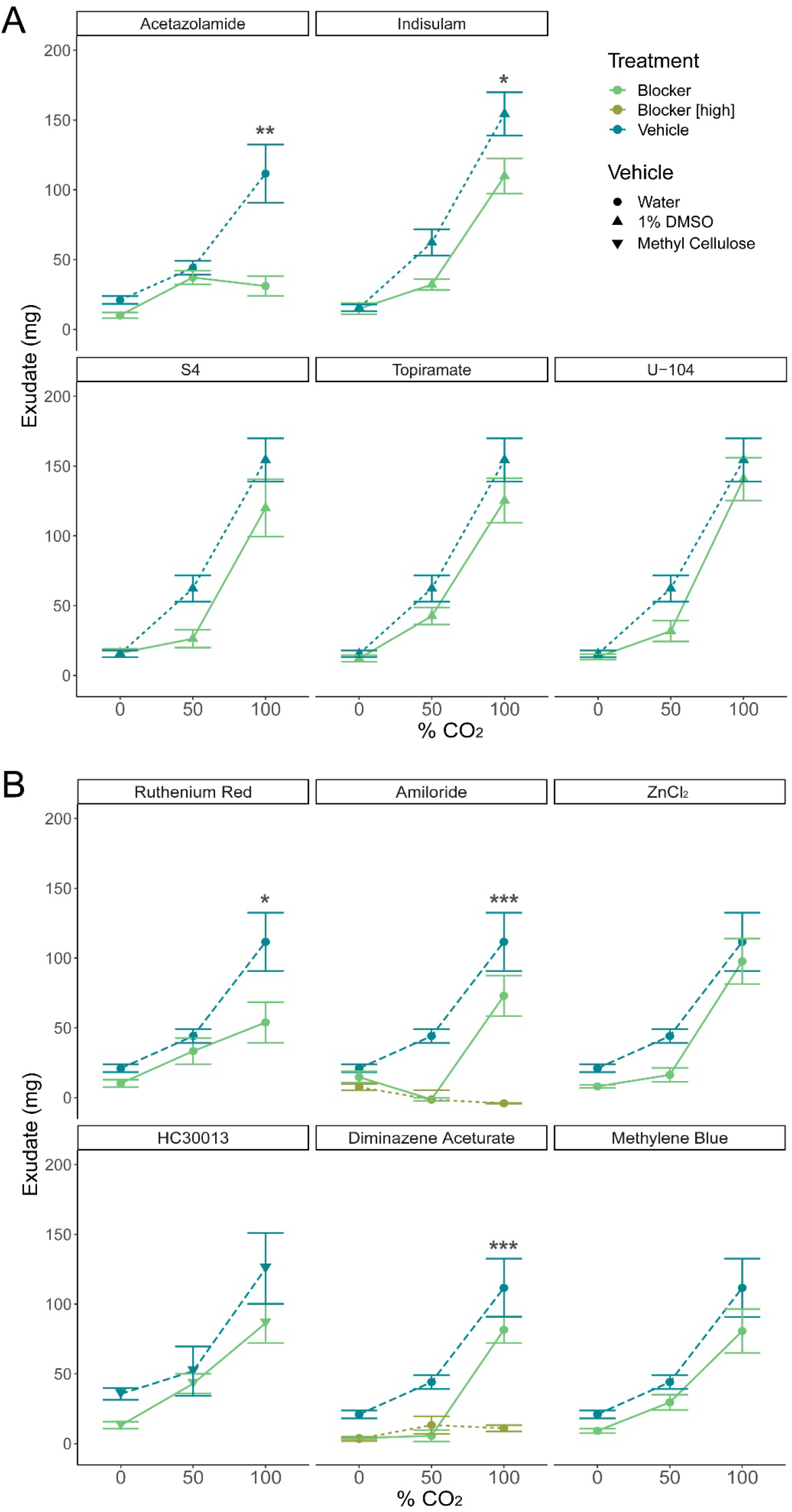
Exudate excretion after treatment with blockers and inhibitors versus treatment with the respective vehicle. A) The carbonic anhydrase inhibitors acetazolamide (general CA inhibitor, p<0.01, n=8-13) and indisulam (CA IX/XII inhibitor, p<0.05, n=6-8) significantly muted the exudate response by a two-way ANOVA and significant differences were detected via Tukey’s HSD test. S4 (CA IX inhibitor, p>0.05, n=5-8), U-104 (CA IX/XII inhibitor, p>0.05, n=6-8) and topiramate (CA II/IV inhibitor, p>0.05, n=6-8) did not significantly alter the response. B) Receptor blockers amiloride (ENaC blocker, p<0.0001, n=3-8), diminazene aceturate (ASIC3 blocker, p<0.0001, n=3-8), and ruthenium red (Ca2+ channel blocker, p<0.01, n=5-13) significantly muted the exudate response by a two-way ANOVA, and significant differences were detected via Tukey’s HSD test. HC030013 (TRPA1 blocker, p>0.05, n=6-12), methylene blue (guanylate cyclase inhibitor, p>0.05, n=6-8), and ZnCl2 (OTOP1 blocker, p>0.05, n=6-8) did not significantly alter the response. Graphed values are means ± SEM; * p <0.05, ** p <0.01, *** p <0.001 via Tukey’s HSD test.

Of the receptor blockers, the broad-spectrum ENaC/ASIC blocker amiloride (p<0.0001, n=3-8) and the ASIC3 blocker diminazene aceturate (p<0.0001, n=3-8), and the Ca^2+^ channel blocker ruthenium red (p<0.05, n=5-13) significantly muted the response to CO_2_ as detected by two-way ANOVA and Tukeys HSD test (Figure 7B). The TRPA1 channel inhibitor HC030013, the guanylate cyclase inhibitor methylene blue, and ZnCl_2_ that blocks OTOP1 channels did not significantly alter exudate production (p>.05, n=5-13).

To control for effects of the blockers on the mechanisms of exudate release rather than detection of CO_2_, the positive control AITC was tested after exposure to acetazolamide, amiloride, diminazene aceturate, indisulam, and HC030013 (Figure 8). Acetazolamide (p>0.05, n=4), 1 mM amiloride (p>.05, n=4), and indisulam (p>0.05, n=5) did not significantly mute the response to AITC by a TukeysHSD test, but HC030013 (p<0.001, n=7), 5 mM amiloride (p<0.001, n=4), 0.05 mM diminazene aceturate (p<0.01, n=5), and 0.1 mM diminazene aceturate (p<0.05, n=5) did. HC030013 is a TRPA1 blocker and amiloride can act as one at higher concentrations (Banke, 2011; Eid et al, 2008) and predictably blocked AITC detection. The vehicle for AITC, mineral oil, produced no significant response on its own when compared to 0% CO2 (p=1.00, n=4). Taken together these results indicate that Acetazolamide, 1 mM amiloride, and indisulam have no detectable effect on the ability of earthworms to release of exudate when confronted with noxious stimuli.

**Figure 8.**
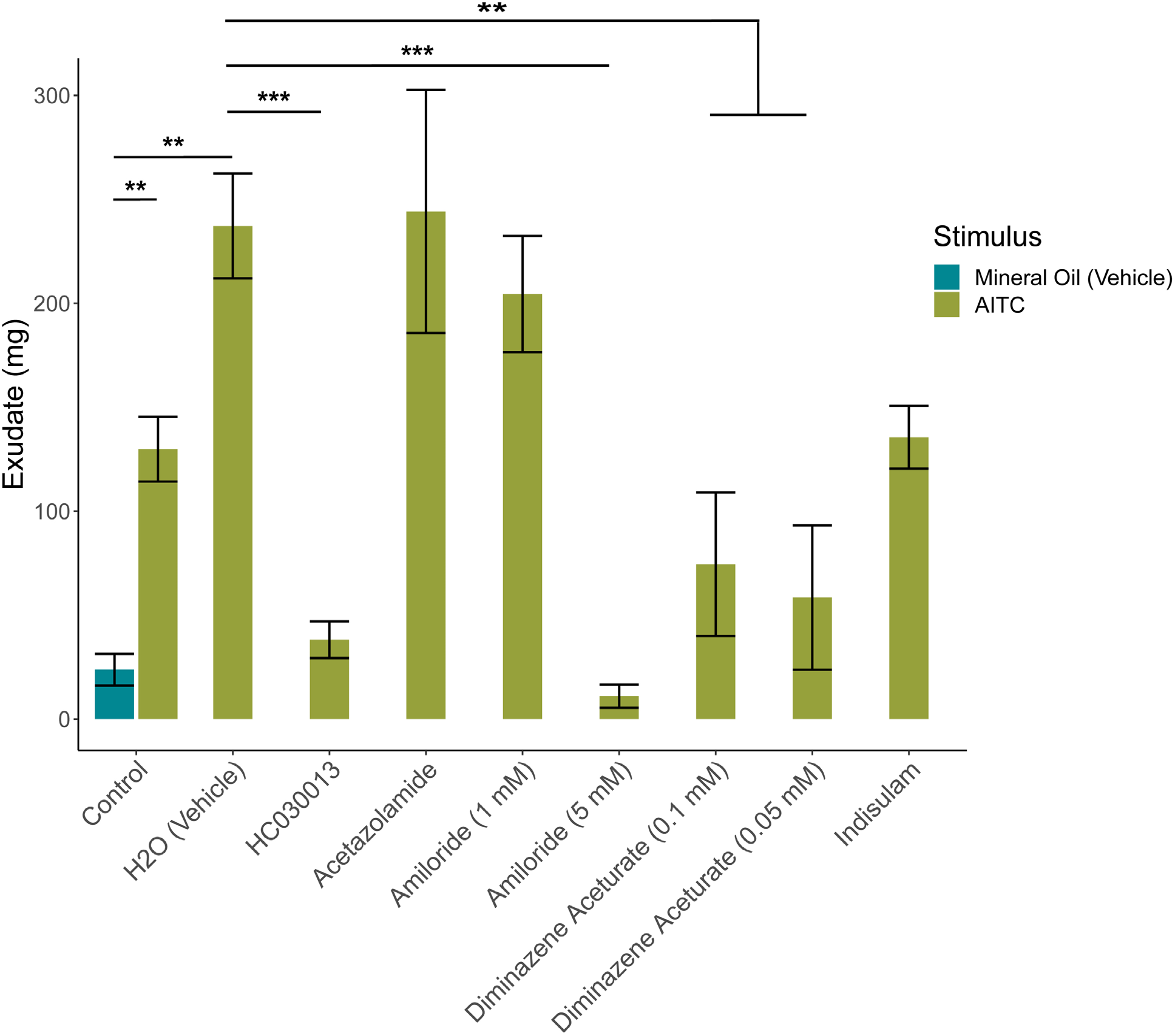
AITC induced exudate excretion after treatment with blockers. AITC significantly increased exudate production (p < 0.01, n=8) and was altered by blocker treatment (p < 0.0001, n=4) compared to control (n=4) via two-way ANOVA. The response to AITC was significantly muted by the TRPA1 blocker HC030013 (p<0.0001, n=7), 5mM amiloride (p<0.0001, n=4), 0.05mM diminazene aceturate (p<0.0001, n=5) and 0.1mM diminazene aceturate (p<001, n=5) via Tukey’s HSD test; while acetazolamide (n=4) and indisulam (n=5) did not significantly alter exudate production. Graphed values are means ± SEM; * p <0.05, ** p <0.01, *** p <0.001.

### Organic Acids

Carbonic acid is a weak acid; like carbonic acid, other weak acids are also known to stimulate nociceptors in animals (Wang et al., 2011). We decided to test whether earthworms were also sensitive to a series of other weak acids. Indeed, they excreted significantly more exudate in response to increasing concentrations of formic acid, acetic acid, and propionic acid (p<0.0001, n=3-8, two-way ANOVA) (Figure 9A). This response was significantly muted in a dosage-dependent manner after treatment with the sodium channel and ASIC blocker amiloride (p<0.0001, n=4-8, two-way ANOVA) (Figure 9B). This result supports the hypothesis that the detection of other weak acids also requires channels from the ENaC/DEG family.

**Figure 9.**
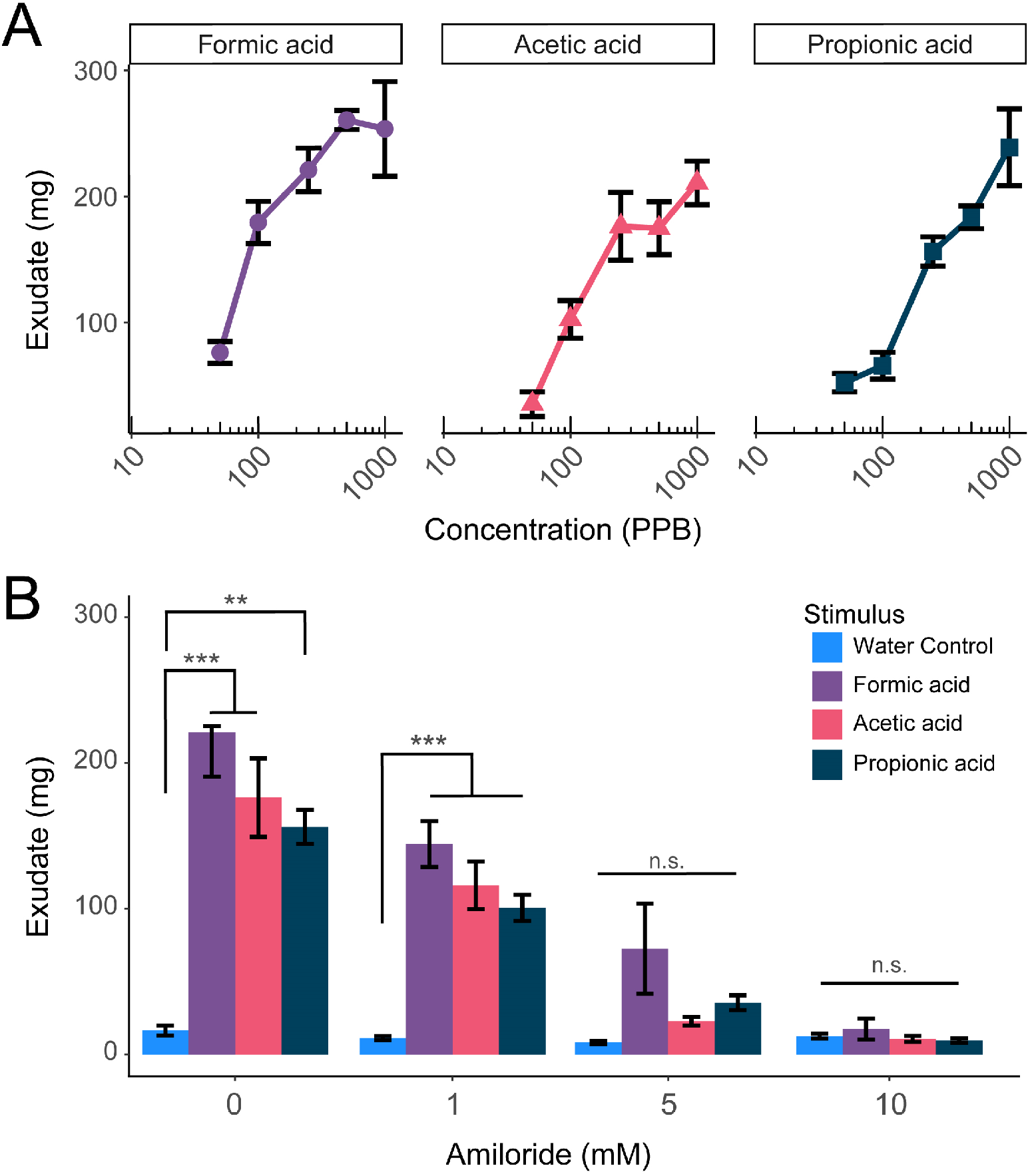
Earthworms produce exudate in response to weak organic acids. (A) Earthworms excrete significantly more exudate in response to increasing concentrations of formic acid, acetic acid, and propionic acid (p<0.0001, two-way ANOVA). (B) This response is significantly muted by amiloride treatment in a dosage-dependent manner (p<0.0001, two-way ANOVA). Graphed values are means ± SEM (n=4-8); * p <0.05, ** p <0.01, *** p <0.001 via Tukey’s HSD test.

## Discussion

This study demonstrates high CO_2_ tolerance in *D. veneta* and provides initial evidence of the molecular mechanisms behind CO_2_ detection in this species, giving insight into how earthworms detect chemicals in their environment and contributing to ongoing investigations into mechanisms of CO_2_ detection across species. The chemosensory capabilities of earthworms were famously discussed by Darwin and have been the subject of anatomical, physiology and most often ecological studies since that time. There is, however, a comparative dearth of publications that examine earthworm biology with an eye towards identifying the molecular mechanisms that mediate their physiology and behavior (Stürzenbaum et al., 2009). In fact, we believe that this study is the first to use modern sequencing technologies to elucidate the molecular mechanism responsible for any type of earthworm chemosensation.

### CO_2_ Aversion and Tolerance

Because earthworms excrete exudate in response to noxious stimuli (Heredia et al., 2008; Rivera et al., 2020), we used the amount of exudate produced as a proxy for aversion in the exudate assay. We found that CO_2_ is noxious to earthworms in a dosage-dependent manner (Figure 1C). Many species find CO_2_ noxious at high concentrations, including rats, nematodes, and fruit flies (Hallem & Sternburg, 2008; Breugel et al., 2018; Améndola & Weary, 2019). High concentrations of CO_2_ also anesthetize many species, including rats and fruit flies (Seiger & Kink, 1993; Danneman & Walshaw, 1997), but we did not observe this in earthworms even with 100% CO_2_. Low concentrations of CO_2_ (ex: 25%) produced no significant response in earthworms, though such concentrations are noxious in other species including rats (25%; Améndola & Weary, 2019), nematodes (10%; Hallem & Sternburg, 2008), and fruit flies (5% when not foraging; Breugel et al, 2018). Concentrations as high as 45% have been used in human irritation studies, suggesting similar tolerance, although in a much larger animal (Shusterman Avila, 2003; Hummel et al., 1998). Earthworms may be resistant to high CO_2_ concentrations due to high exposure in their environment, with mechanistic adaptations underlying such resistance.

Increasing CO_2_ and therefore increasing carbonic acid is usually considered a proxy measure for hypoxia. There is a precedent for subterranean organisms to be more tolerant of CO_2_ because of their burrows being hypoxic environments. Naked-mole rats have a proton-insensitive ASIC3 channel among other adaptations to their high-CO_2_ environment (Schuhmacher et al, 2018); earthworms may have similar adaptations. While subterranean CO_2_ levels are estimated to be lower than what earthworms are responding to in this study (0.04% to 13.0%), those estimates vary widely (Amundson & Davidson, 1990; Scott, 2011). Additionally, earthworms are often observed to aggregate into balls of 6 to 12 inches in diameter containing hundreds of individuals (Gates, 1961). CO_2_ levels in a subterranean ball of earthworms may exceed the general estimates of subterranean CO_2_ in soil; we are not aware of any studies that attempt to model or measure CO_2_ levels found in aggregates of earthworms. However, we do expect a writhing mass of metabolizing animal tissue to constantly consume oxygen and generate CO_2_ that would not easily diffuse in a subterranean environment. Perhaps together, earthworms produce CO_2_ levels consistent with those tested here and CO_2_ might even be an important chemical signal that limits the size of earthworm aggregates.

Earthworms feed on soil, gaining nutrients from decaying organic matter, bacteria, nematodes, and other microfauna (Curry & Schmit, 2007), many of which emit CO_2_. Low concentrations of CO_2_ may also be appetitive rather than aversive; this is true of mosquitoes and, when in a foraging state, fruit flies (Breugel et al., 2018; Spanoudis et al., 2020). Low concentrations may also use different mechanisms than aversive high concentrations. Insects like mosquitoes and fruit flies, however, detect CO_2_ through ionotropic gustatory receptors, which we found no evidence of in the *D. veneta* transcriptome.

### CO_2_ Adaptations

Earthworm mucus, which coats the epithelium under basal conditions, keeps the skin moist for respiration, aids with locomotion, and acts as a buffer (Zhang et al, 2016; Schrader et al., 1994). This mucus consists of mucin proteins secreted by epithelial cells, mainly goblet cells (Cunha et al., 2011; Rubin, 2014). While little is known about the physiology of earthworm goblet cells, they are well-characterized in the vertebrate respiratory epithelium, which earthworm epithelium resembles histologically (Cunha et al, 2011) and on the ultrastructual level (Sundararaman & Gupta, 1992). If the physiology of goblet cells is as highly conserved between the two tissues as the anatomy, then mucin secretion is likely SNARE-complex dependent exocytosis (Adler et al., 2013), with mucus thickness influenced by the ions transported out of epithelial cells, particularly Cl^-^ (Rogers, 2007).

Exudate, which we define here as the substance excreted in response to noxious stimuli beyond the basal mucus coating, acts as an alarm signal to other earthworms and is attractive to some predators, such as garter snakes (Jiang et al., 1990). Protein and carbohydrate-rich mucus vesicles containing auto-fluorescent membrane-bound chloragosome granules are excreted from between the earthworm’s segments and lubricated by coelomic fluid (Roots and Johnston, 1966; Heredia et al., 2008; Zhang et al., 2016; Guhra et al., 2020). Studies in the closely related *Eisenia fetida* show that the vesicles form strands outside the body, which differs from the globules we observed in *D. veneta* (Heredia et al., 2008). Exudate chemical composition also differs between earthworm species, including in *D. veneta* α-nicotinamide riboside and the organic acids fumarate, succinate, malate, and α-ketoglutarate (Bundy et al., 2001; Rochfort et al., 2017).

It is unclear to what extent mucus and exudate are the same. Many studies electrically or chemically stimulate earthworms to release exudate in order to collect and study mucus (Jiang et al., 1990; Zhang et al., 2016), and *E. fetida* excretes small amounts of exudate proteins under basal conditions (Rivera et al, 2020; Heredia et al, 2008). The most recent research in the field has shown that exudate produced in response to varying intensities of electrical stimuli differ in pH and concentration of amino acids, nitrogen, potassium, and phosphorus (Huan et al, 2023). Our transcriptome data suggest that homologs to secreted CAVI may be one component of both exudate and mucus (Figure 2, S1, transcript ID: 67832_c1_g1_i5 & 76341_c1_g1_i1, 76341_c1_g1_i17). CAVI or gustin is secreted in saliva and airway surface liquid in mammals and is known to play a role in chemosensory perception (Leinonen et al., 2004; Fábián et al., 2015).

In our study, earthworms rinsed with deionized water produced more exudate in response to CO_2_ compared to untreated worms (Figure 1C). We hypothesize that the water washed off the earthworms’ mucus. A scant number of studies have thoroughly examined or compared the composition and the chemical properties of earthworm mucus and exudate. We predict, however, that mucus may have neutralized the low-pH environment created from CO_2_ and water by carbonic anhydrase. Rinsing the mucus away with deionized water likely allowed more CO_2_ to dissolve and be converted to bicarbonate and protons by carbonic anhydrase, producing more protons to activate receptors. Alternatively, mucus may have simply physically blocked the epithelium from irritants. That treatment with water also increased the aversive response to AITC supports this hypothesis but does not preclude both mechanisms from contributing to the observed response. These competing hypotheses emphasize the need for more research focusing on the molecular mechanisms that underlie earthworm chemical senses.

### Earthworm Carbonic Anhydrases

While we acknowledge that these concentrations of CO_2_ are much higher than what an earthworm is likely to encounter in its subterranean environment (up to 13% CO_2_) (Amundson & Davidson, 1990), the “non-ecological” concentrations used in this study allowed for investigation of the mechanisms behind CO_2_ aversion. Repetition of the exudate assay after exposure to blockers demonstrated that carbonic anhydrase is necessary for aversion to high concentrations of CO_2_, as the general carbonic anhydrase inhibitor acetazolamide and the carbonic anhydrase IX/XII inhibitor indisulam both significantly muted responses to CO_2_ (Figure 7A). Earthworms thus likely have functioning carbonic anhydrases and may have CAIX and CAXII orthologues. That the CAIX/XII inhibitor U-104 and CAIX inhibitor S4 had little effect on CO_2_-induced irritation, however, may contradict this theory or suggest that the CAIX/XII orthologues have diverged such that these blockers are no longer effective at inhibiting the enzyme or that the isoforms more closely resemble CAIX.

In mammals, CAXII, CAII, CAVB, and CAIV are found in the human nasal mucosa (Tarun et al., 2003), and CAIV is required for sour taste responses to carbon dioxide (Chandrashekar et al., 2009). The CAII/IV inhibitor topiramate had no significant effect in our study, suggesting that CAII and CAIV orthologues are either not involved in earthworm CO_2_ detection, or are so divergent from those of organisms on which topiramate has been tested that the blocker simply does not have an effect. CAVI, one of the only known secreted isoforms, is found in mammalian saliva and is associated with taste (Henkin et al, 1999; Fábián et al., 2015). We notably found CAVI orthologues in the *D. veneta* epithelial transcriptome but did not have access to a CAVI inhibitor for pharmacological studies. Since CAVI is secreted protein, it is likely that rinsing the worms with water removed any CAVI present in the epithelial surface, raising the possibility that it might be responsible for the differential response seen before and after rising (Figure 1C). It is nonetheless likely that earthworms secrete a CAVI orthologue in their mucus or exudate that plays some role in CO_2_ or acid detection.

Multiple carbonic anhydrase isoforms may contribute to earthworms’ response to CO_2_; this would explain the larger effect by the general carbonic anhydrase inhibitor acetazolamide compared to the inhibitors of single isoforms. Additionally, given the relatively low percentage identity of earthworm CA to its homologs, that acetazolamide retained its effectiveness in muting the earthworms response to CO_2_ validates it at a broad-spectrum CA inhibitor. The biological redundancy observed in earthworms response to CO_2_ is likely indicative of the general importance of CO_2_ detection to all animals, or could be evidence that carbonic anhydrases are of particular importance to the survivability of earthworms. That carbonic anhydrase is required for a response to CO_2_ also suggests that earthworms’ sensory receptors detect protons or bicarbonate rather than CO_2_.

### Mechanisms of CO_2_ Detection

Earthworm sensory cells are found on their epithelium, either alone as solitary chemoreceptor cells (SCCs) or grouped into epithelial sensory organs (ESOs), which all project to a ventral nerve cord (Hess, 1925; Csoknya et al., 2005; Kizsler et al., 2012). Receptors activated by CO_2_, bicarbonate, or protons are likely found on the membranes of these cells. There is evidence of epithelial sodium channels including acid-sensing ion channels (Figure 3), receptor-type guanylate cyclases (Figure 4), otopetrins (Figure 5), and TRPA1s (Figure 6) in the *D. veneta* transcriptome. Guanylate cyclases, OTOP channels, and TRPA1 channels are likely not required for CO_2_ detection in *D. veneta*, as their blockers did not significantly affect earthworms’ responses to CO_2_ (Figure 7B). Because carbonic anhydrase is required for CO_2_ aversion (Figure 7A) and there were no matches for the gustatory receptor family that detects CO_2_ in arthropods, it is also unlikely that CO_2_-specific sensory cells are required.

Amiloride, however, did mute the exudate response to CO_2_, suggesting that an epithelial sodium channel (ENaC) is required for CO_2_ detection and aversion (Figure 7B). The ASIC3 blocker diminazene aceturate also muted responses to CO_2_, but the role of ASIC3 cannot be confirmed because it also muted the response to the positive control AITC, suggesting that it may impact the mechanism of exudate excretion rather than CO_2_ detection (Figure 8). High concentrations of amiloride also muted responses to AITC, but, because the ENaC blocker amiloride is also a TRPA1 blocker at high concentrations (Banke, 2011), this response was somewhat expected. Low concentrations of amiloride muted responses to CO_2_ but not AITC, further suggesting the role of the ENaC superfamily in CO_2_ detection.

This is corroborated by the responses to organic acids, which were also muted by amiloride (Figure 8), suggesting that the same mechanism is responsible for acid detection and CO_2_ detection. Formic acid, acetic acid, and propionic acid are all weak acids like carbonic acid. That amiloride muted the response to these three organic acids as well as to CO_2_ provides additional evidence that the mechanism of CO_2_ detection in *D. veneta* detects a product of the carbonic anhydrase reaction (carbonic acid, protons, or bicarbonate; Figure 1A) and that an ASIC homolog is required for this response.

### Future Research and Conclusions

Earthworms are critical to many terrestrial ecosystems, yet we know little about how they make decisions about chemicals in their environment. Additional genomic research in *D. veneta* and related earthworm species is a necessary next step to understanding earthworm chemical ecology, and high throughput sequencing of *D. veneta* RNA reported here provides the opportunity to do so. RNAi or CRISPR studies are the logical next step to corroborate the pharmacological evidence for the roles of carbonic anhydrase and ENaCs in earthworm CO_2_ detection presented in this study. These methods, however, are entirely dependent on discovering a reliable technique for collecting very early-stage earthworm embryos, and we have yet to identify any protocols to do so in the literature—another datum highlighting the need to focus more basic cellular biology research on these organisms. Phylogenetic analyses of chemosensory genes across annelids will continue to provide insight into patterns of chemosensory evolution, potentially revealing adaptations to earthworms’ low-light, high-CO_2_ environment.

CO_2_ is a critical signaling molecule that earthworms encounter at high concentrations. This research could thus contribute to a necessary understanding of what attracts and repels earthworms, which has potential applications in agriculture and invasive species management. Earthworms have critical roles in soil fertility, and this research may consequently have additional implications in ecosystem health and biodiversity. Research on responses to CO_2_ may be of particular importance in the context of climate change: as CO_2_ concentrations in the atmosphere increase, CO_2_ concentrations in the soil may increase as well. These results suggest that increasing concentrations of CO_2_ in the soil won’t be aversive to earthworms, demonstrating their potential resilience to a changing environment. There are also potential applications in the assessment and treatment of acidic soil, as our data supports previous studies describing earthworm mucus as an effective buffer (Zhang et al, 2016; Schrader et al, 1994).

These data may also provide insight into CO_2_ detection and chemosensation across species. Because earthworms respire (e.g. “breathe”) through their skin, their entire body is a respiratory epithelium (Karaca, 2011). This makes them an excellent model for chemosensation in a respiratory epithelium across invertebrate and vertebrate species, including CO_2_ and acid detection. Our investigation of this topic has contributed to our understanding of the mechanisms earthworms use to detect carbon dioxide and provides yet another example of how these molecular mechanisms are highly conserved among metazoa.

## Supplementary material

S1: SmithEtal2023-candidates.xlsx - Excel file containing full Trinotate annotations for candidate transcripts in this study.

S2: SmithEtal2023-Deseq-trinotate-p05.xlsx – Excel file containing two worksheets. One contains the significant results from differential sequencing analysis between prostomium and mid-segment; the other worksheet contains the trinotate annotations for those transcripts. None of the significantly differentially expressed transcripts are implicated in CO2 or weak acid detection.

## Supporting information

S1 SmithEtal2023-canidates

S2: SmithEtal2023-Deseq-trinotate-p05.xlsx

## Data availability statements

The RNA-sequencing datasets generated for this study can be found in the NIH’s National Library of Medicine’s National Center for Biotechnology Information’s Sequence Read Archive (https://www.ncbi.nlm.nih.gov/sra). The Bioproject ID number for this data set is PRJNA1026068. The raw exudate data and scripts used to analyze both the RNA-sequencing and exudate assay data can be found in the following GitHub repository: https://github.com/JakeSaunders/Smith_etal_2023_Earthworm_CO2

## Ethics Statement

Ethical review and approval were not required for studies on annelids in accordance with the local legislation and institutional requirements.

## Acknowledgments

Computations for this study were performed using the Wake Forest University (WFU) High-Performance Computing Facility, a centrally managed computational resource available to WFU researchers including faculty, staff, students, and collaborators (Information Systems and Wake Forest University, 2021). Some data and text included in this manuscript originally appeared in ES’s master’s thesis (Smith, 2021).

